# Effects of cross-generational inbreeding and *Wolbachia* infection on sex ratio and life-history traits in parthenium beetle

**DOI:** 10.64898/2026.05.08.723828

**Authors:** Biswajita Swain, Ranjit Kumar Sahoo

## Abstract

Sex ratio is a key demographic parameter shaping population dynamics and evolutionary trajectories. In biocontrol agents, demographic bottlenecks during species introduction to a new habitat and subsequent mass rearing can elevate inbreeding, potentially biasing sex ratios through sex-specific mortality associated with inbreeding depression. Moreover, reproductive endosymbionts such as *Wolbachia* are known to manipulate host reproduction and further skew sex ratios. However, the relative contributions of these processes to sex-ratio variation remain poorly resolved. In this study, we evaluated the effects of cross-generational full-sibling inbreeding and *Wolbachia* infection on sex ratio and key life-history traits in the biocontrol beetle *Zygogramma bicolorata* using controlled laboratory crosses across three generations. Inbreeding did not significantly alter offspring sex ratio, which remained close to parity across generations, while pupal mortality increased in later generations, consistent with delayed expression of inbreeding depression. Adult body weight remained largely unaffected by inbreeding. *Wolbachia* infection was detected in a subset of females and was associated with a modest but significant increase in female-biased offspring production, although the effect was variable across lineages. Strain typing identified a single supergroup A *Wolbachia*, consistent with previous descriptions of the *w*Bic strain from this species. These findings indicate that sex-ratio variation in introduced populations of *Z. bicolorata* is not driven by inbreeding alone but instead emerges from the interaction between demographic processes and symbiont-mediated effects, providing crucial insights for optimizing biocontrol programs where sex-ratio stability is essential for population establishment and persistence.

**Significance:** Sex ratio is a key determinant of population growth and stability – the essential parameters determining success of biocontrol programs. Yet, the mechanisms shaping sex-ratio variation remain poorly resolved. Using controlled crosses in *Zygogramma bicolorata*, we show that short-term inbreeding does not directly alter sex allocation, despite inducing delayed fitness costs through increased pupal mortality. In contrast, *Wolbachia* infection contributes to female-biased offspring production, although with variable outcome across lineages. These findings demonstrate that sex-ratio variation in *Z. bicolorata* arises from the interaction of demographic processes and symbiont effects, rather than a single mechanism, with important implications for predicting the establishment, persistence, and efficacy of mass-reared biocontrol populations.

## Introduction

In sexually reproducing dioecious organisms, population sex ratio is a key demographic parameter shaping both short-term population growth and long-term evolutionary trajectories. By structuring mating opportunities and reproductive output, sex ratio can influence mating system dynamics and the distribution of allelic variation within populations. Persistent deviations from an expected equilibrium can, under certain conditions, accelerate allelic fixation or loss, thereby reducing genetic diversity (Ciftci etal., 2020). Empirical studies have demonstrated such links, for example, in *Antennaria dioica*, population sex ratio – rather than population size – was shown to correlate with levels of genetic diversity (Rosche et al., 2018).

However, patterns of sex-ratio variation observed across taxa indicate that the underlying proximate drivers are multifactorial (Dubey and Singh, 2022). Classical theory predicts that life-history traits such as mating system, dispersal, and sex allocation strategies can modulate sex ratios, while ecological factors – including population structure and resource distribution – further shape these outcomes. Population inbreeding is also predicted to impact sex ratios in a species-specific manner, for example, in hymenopteran species, inbreeding lead to higher diploid male offspring (Zaviezo et al., 2017; Zhou et al., 2007). Yet, theoretical and empirical studies suggest that the effect of inbreeding on sex ratio is often weak and likely manifests through the sex-specific outcome of inbreeding depression (Robinson et al., 2014; Frankham and Wilcken, 2006). Furthermore, maternally inherited endosymbionts such as *Wolbachia* are well known to distort host reproduction through mechanisms such as feminization, male killing, or cytoplasmic incompatibility, thereby generating biased sex ratios in natural populations (Werren et al., 2008). Yet, the relative contribution of these factors remains poorly understood.

In this study, we examine some of these processes for their effects on sex ratio and other key life-history traits in the leaf-feeding beetle *Zygogramma bicolorata* (recently proposed to rename it as *Calligrapha bicolorata* (Shawn et al., 2024)), a biological control agent of the invasive weed *Parthenium hysterophorus*. Native to the Americas, *Z. bicolorata* has been widely introduced across Asia, Africa, and Australia following protocol-driven biocontrol programs (Dhileepan and Strathie 2009; Dhileepan and Wilmot Senaratne 2009; CABI 2021). Such programs often involve strong demographic bottlenecks and periods of controlled mass rearing, both of which can impose varying degrees of inbreeding. In India, for instance, the founding population was established from a small number of individuals that were subsequently propagated under laboratory conditions prior to release (Mishra, 2012; Sushilkumar, 2009; Jayanth, 1987; Jayanth and Nagarkatti, 1987). Lab and field observations have consistently reported a female-biased sex ratio across introduced populations (Sushilkumar et al., 2022; Hasan & Ansari 2016; Siddhapara et al., 2012; Jayanth & Bali 1993), yet the underlying causes of this bias remain unresolved. Here, using controlled breeding experiments combined with molecular screening, we evaluate two non-mutually exclusive mechanisms that could shape sex-ratio variation in this species: cross-generational inbreeding and infection by *Wolbachia*. In addition, we assess the impact of inbreeding on key life-history traits, including survival and body size, to determine whether potential fitness costs are decoupled from sex-ratio dynamics.

## Methods

### Study species

For our current investigation, we sampled beetle females from the surrounding area of the city of Hyderabad, India to establish the laboratory populations. Beetles were collected from the same location (17.346065N, 78.404334E) at two distinct time periods forming two independent experimental setups. Therefore, our study has two batches of laboratory populations: one started from eleven females collected during December 2021 and another from six females collected during October 2022. In the laboratory, the beetle populations were reared on both field-collected and nursery-grown *P. hysterophorus* leaves. Populations of both batches were maintained at ambient room temperature, following the protocol described below.

### Laboratory maintenance protocol

Wild collected females were placed individually in 2.5L (21*15*8 cm^3^) aerated plastic containers along with the fresh leaves of *P. hysterophorus* base-wrapped with moist paper towel. After 48 hrs, eggs were collected from the containers. On the third and fourth days of egg collection, larvae were sampled and placed in petri dishes (150*20 mm^2^). Each petri dish had a maximum of ten larvae supplied with a base-wrapped fresh leaf. Leaf was replaced alternate days. On the seventh day of larval collection, to prevent crowding and maintain hygiene, the maximum number of larvae in each petri dish was reduced to five by adding more petri dishes to the maintenance protocol.

Towards the twelfth day of larval collection, fourth instar larvae at pre-pupal stage were supplied with 5% moistened sand for pupation. Each 250 ml pupation box (dia=8 cm, ht=8 cm) was filled with the moistened sand up to 5cm height and served as the pupation bed for maximum five individuals. Moistened sand was prepared by adding required amount of tap water to the autoclaved and subsequently sun-dried sand. From the 10^th^ day of pupation, adults started eclosing.

Freshly eclosed adults were sexed within 24 hrs of eclosion (as described in the next section) and placed in the plastic container of 2.5L capacity with a maximum of 15 individuals each. Males and females were kept in separate boxes. Sufficient fresh leaves were supplied on alternate days. Ten days old adults from both the sexes were placed together in mating chambers for 48 hrs. Females were then separated and kept individually in petri dishes (150*20 mm^2^) supplied with fresh leaves. After 48 hrs of egg laying window, eggs were collected for the next generation.

### Sexual dimorphism

The beetle *Z. bicolorata* exhibits a polygamy mating system where both male and female potentially mate with two or more members of the opposite sex. The male and female sexes are distinguishable by the dimorphism in the last abdominal sternite, which is slightly serrated in case of male and entire in female (Jayanth 1991; Jayanth & Bali 1993). Faint depression at the center of the last abdominal sternite is another recognizable feature in male (Jayanth 1991; McClay, 1980). Nevertheless, we used the presence of serrated or straight terminal segment as the characteristic feature to distinguish male and female, prominently. Presence/absence of faint depression was used as a confirmatory character in the cases of ambiguity.

### Experimental design

The adult offspring (Generation F_1_) of wild collected females (Generation F_0_) were mated strategically on one-to-one basis to facilitate monitoring their pedigree. In each case, the mating pair had 48 hrs of mating window followed by 48 hrs of egg-laying for the mated female. Offspring of each mated pair were reared separately. In the next generation (Generation F_2_), the adults underwent either (i) no-sibling mating where the mating partners didn’t share the parents, or (ii) full-sibling mating where the partners share both the parents. The former no-sibling mating types represented the ‘random’ category and the latter full-sibling ones represented the ‘inbreeding’ category. In the inbreeding category, full-sibling mating was conducted for two or three subsequent generations depending upon the mating partner availability and mating success. Throughout the experiment, we followed the rearing protocol as described above.

### Test for *Wolbachia* infection

All beetle samples from the breeding experiment were preserved individually in deep freezer at -80º C. Required samples were retrieved from freezer and processed for DNA extraction. We used tissue from the legs and followed the protocol that has been adapted earlier for the extraction of high molecular weight genomic DNA in this beetle (Sahoo et al., 2023) and in other species (Dias et al. 2021; Yang et al. 2021; Miller et al., 1988). A brief description has been given below.

Briefly, we homogenized tissue sample from each individual in 600 µl lysis buffer (10mM Tris-HCL, 100 mM EDTA, 100 mM NaCl). Homogenized sample was supplied with 40 µl of 10% SDS and 100 µl proteinase K solution (1 mg/ml proteinase K, 1% SDS, 4 mM EDTA), and allowed for overnight digestion at 37º C. Subsequently, 240 µl of 5 M NaCl was added to neutralize the nucleic acid and separate it from other undesired components through centrifugation. The collected supernatant was further supplied with 1 ml pure ethanol for DNA precipitation. Subsequently, the DNA pellet was air dried and suspended in 50-100 µl of sterilized water.

DNA extraction quality was first assessed by amplifying the mitochondrial gene cytochrome oxidase I (COI: primer pair LCO/HCO) (Wahlberg and Wheat, 2008). Samples positive for COI were tested for *Wolbachia* infection by amplifying two *Wolbachia-*specific gene regions, *Wolbachia* surface protein gene *wsp* (primer pair: wsp 81F/wsp 691R) (Zhou et al., 1998) and cell cycle protein gene *ftsZ* (primer pair:ftsZ_F1/ftsZ_R1) (Baldo et al., 2006). Samples positive for both *wsp* and *ftsZ* were assigned positive for *Wolbachia* infection.

### *Wolbachia* Strain characterization

To characterize the *Wolbachia* strain type in our samples, we sequenced four additional gene loci (*fbpA, hcpA, gatB, coxA*) for five selected positive samples. Subsequently, we used five loci (*ftsZ, fbpA, hcpA, gatB, coxA*) for multi-locus sequence typing (MLST) and one locus (*wsp*) for WSP-typing (Baldo et al., 2006) of these samples. First, we manually corrected the sequences, and then, checked each locus for sequence divergence within our samples by aligning them using ClustalW (Thompson et al., 1994) in default mode as implemented in the program MEGA v11 (Tamura et al., 2021). Thereafter, we end-trimmed the sequences as per their lengths from the PubMLST database and searched all sequences for their closest match against the PubMLST *Wolbachia* database (Jolley et al., 2018) and assigned the sequence typings.

To assign a supergroup classification to our strain, we collected 15 *Wolbachia* strains from the PubMLST database for which all five MLST loci were available. The collected data, which covered the strains from A, B, D and F supergroups, were combined with our sequences from five beetle samples to build a phylogeny-based strain grouping. We reconstructed the maximum likelihood (ML) tree using the concatenated MLST loci under the Tamura-Nei (G+I) model with 1000 bootstraps in the program MEGA v11 (Tamura et al., 2021).

## Results

### Effect of inbreeding on sex ratio

Our dataset comprised offspring sex ratio data from 39 mating pairs (with a minimum of three offspring per pair (Fig. S1)) (Supplemental data). Among these, 10 pairs belong to the random mating category, and 29 pairs to the full-sibling inbreeding category, or simply, inbreeding category. Out of 29 inbreeding pairs, 10, 12 and 07 pairs represented first, second and third generation, respectively. We tested whether the cross-generational inbreeding affects the offspring sex ratio in this beetle species. Our analysis showed that inbreeding did not induce any significant change to the median sex ratio across three generations (Kruskal-Wallis (KW) test: chi-squared=2.47, df=2, p-value=0.289) (Fig. 1A). However, when data from the random mating category was added to the analysis, we observed a near significant effect of breeding pairs on the offspring sex ratio (KW test: chi-squared=7.36, df=3, p-value=0.061) (Fig. 1A). This suggests that the offspring sex ratio from random mating pairs is slightly different from that of the inbreeding categories. Further analyses revealed that while the median sex ratio from inbreeding remained close to 1: 1 across generations (Wilcox test against µ=0.5: p >0.1), in the random category, the median sex ratio appeared female-biased at approx. 1.4: 1 (female: male) (Wilcox test against µ=0.5: w=41, p=0.03). Batch-specific analyses of the data concurred these results (Fig. 1B), except for the second generation inbreeding from the batch2. In this particular case, the sex ratio was male-biased at 0.7: 1 (female: male) (Wilcox test against µ=0.5: p=0.06), and differed significantly from the random category in the same batch (Dunn’s test: p=0.026) (Fig. 1B).

**Figure 1.**
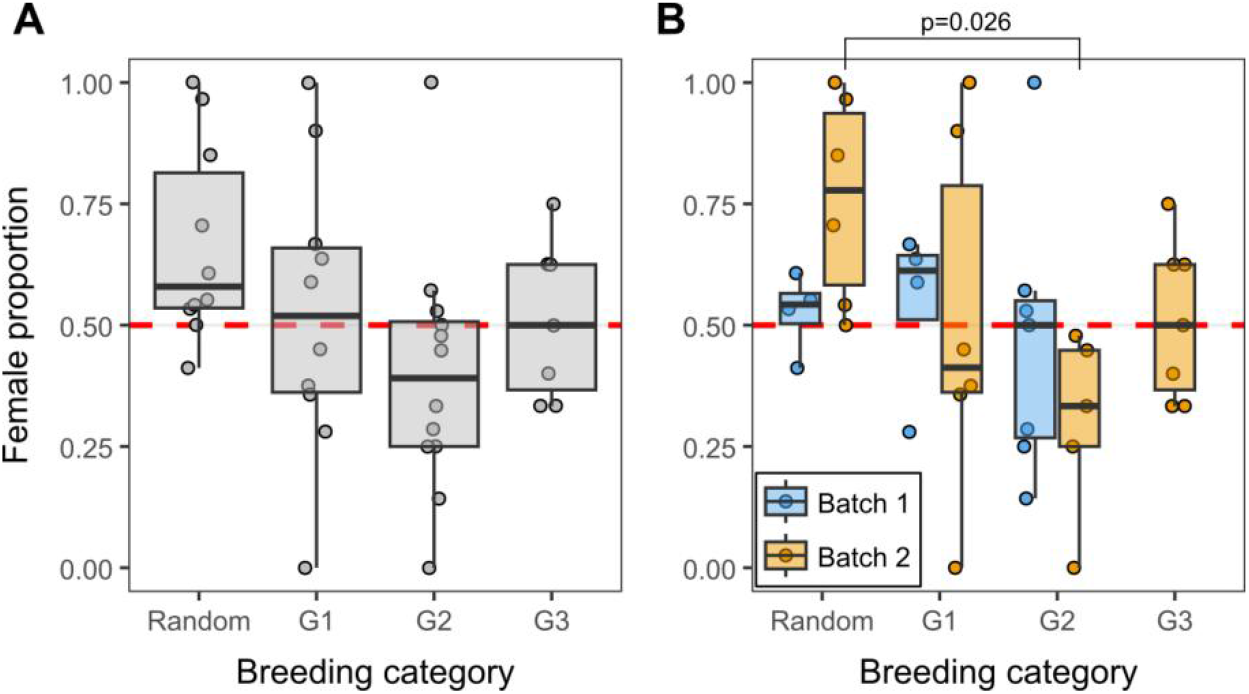
Effect of inbreeding on sex ratio. (A) Box plots showing the proportion of female offspring under random mating and successive generations of inbreeding (G1–G3). Points represent individual mating pairs. The horizontal dashed red line indicates an expected female proportion of 0.5. (B) Batch-wise distribution of the same data. Statistically significant comparisons between groups are indicated with corresponding p-values.

Overall, our dataset suggests no immediate effect of cross-generational inbreeding on the beetle sex ratio within three generations. However, there was a batch-specific effect on the extent of variation in sex ratio across breeding categories. We measured the extent of variation using the robust coefficient of variation (RCV), which is calculated by the ratio between standardized median absolute deviation (MAD) and the median (formula: 1.4826*MAD/median), following Arachchige et al (2020). RCV is a robust measure of relative dispersion of data points in a skewed distribution. Using this measurement, our results showed that, in the random category, the observed sex ratio in the batch2 has relatively higher RCV compared to the batch1 (RCV: batch2=0.579, batch1=0.149) (Table S1). This suggests, mating pairs in batch2 vary widely in their offspring sex ratio compared to that of batch1. Because the two batches differ in their founding members, we anticipate that the initial genetic background is the potential associate of the batch-specific effect. However, this difference was not consistent in the inbreeding category and across generations. For instance, while in batch2, RCV of offspring sex ratio gradually reduced as a consequence of cross-generational inbreeding, it rather increased in the batch1 (Fig. 1B; Table S1).

### Effect of inbreeding on pupal mortality and adult body weight

Next, we checked the effect of inbreeding on other key life-history traits in the beetle species. First, we tested whether cross-generational inbreeding influenced the pupal survival rate. We measured pupal mortality as the ratio of the number of uneclosed pupae to the total number of pupae. We found that the cross-generational inbreeding had significant effect on the pupal mortality (KW test: chi-squared=16.548, df=3, p-value=0.0008) (Fig. 2A), which was primarily because of significantly higher pupal mortality observed in the third generation inbreeding in the batch2 (Dunn’s test: p<0.05). Pupal mortality during the first and second generation inbreeding didn’t vary from each other (Dunn’s test: p-value=1.0) and neither each of them from the random category (Dunn’s test: p-value >0.24). Batch-wise analyses concur these results of pupal mortality pattern for first and second generation inbreeding (Fig. 2B). Nevertheless, we could not detect any potential effect of pupal mortality on the adult sex-ratio across the mating pairs (Spearman correlation Test: H_o_= no association, ρ= -0.169, p=0.304) (Fig. S2).

**Figure 2.**
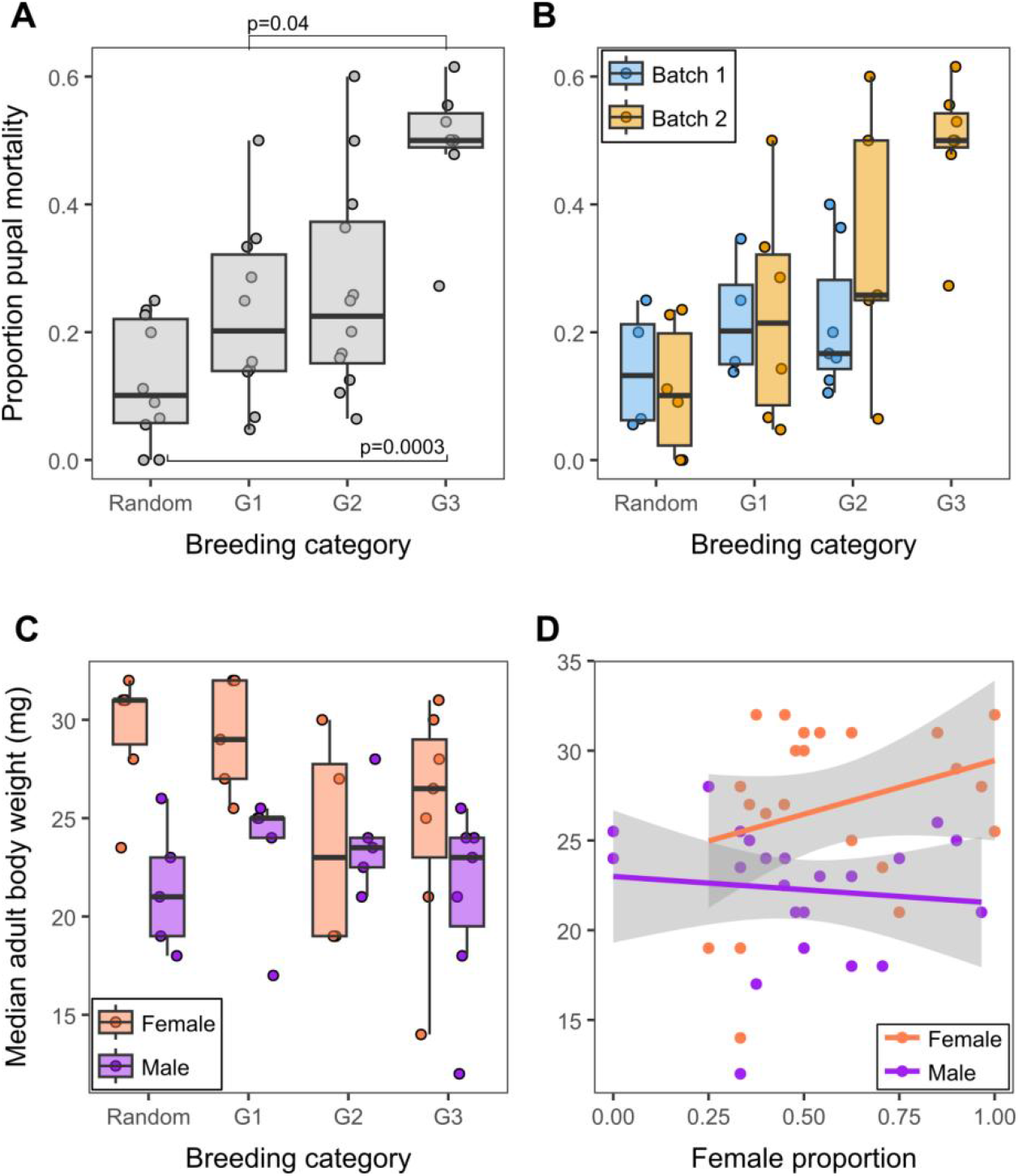
Effects of inbreeding on pupal mortality and adult body weight. (A) Box plots showing the proportion of pupal mortality under random mating and successive generations of inbreeding (G1–G3). Points represent individual mating pairs. Statistical comparisons between groups are indicated with corresponding p-values. (B) Batch-wise distribution of the data shown in (A). (C) Box plots showing median adult body weight under random mating and successive generations of inbreeding (G1–G3) in Batch 2 samples only. Points represent individual mating pairs. (D) Scatter plot illustrating the relationship between female proportion and adult body weight, shown separately for male and female individuals. Points represent individual mating pairs. Solid lines represent linear model fits (R function ‘lm’), with shaded areas indicating 95% confidence intervals.

Next, we tested the inbreeding effect on the adult body weight distribution, and hypothesized that cross-generational inbreeding would result in reduction of offspring body weight across generations. We measured the whole body weight of freshly eclosed adults within 24 hours of eclosion and prior to any food ingestion. Our results, from batch2 samples only, show that cross-generational inbreeding has no apparent effect on the adult body weight distribution, in both male (KW test: chi-squared=1.568, df=2, p=0.457) and female (KW test: chi-squared=2.678, df=2, p=0.262) individuals (Fig. 2C). Distribution in the inbreeding classes also did not differ from the random breeding category (KW test: p>0.136) (Fig. 2C). Notably, in the random breeding category, the median female body weight (31.00 mg) is ∼1.48 times higher than that of the males (21.00 mg). As a consequence of inbreeding, however, the difference between the female and male body weight diminished, where female to male weight ratio was 1.16 (29/25), 0.98 (23/23.5) and 1.15 (26.5/23), respectively for first to third generation inbreeding. We observed no general pattern in variation of the adult body weight, in both male and female, in relation to the offspring sex-ratio (Spearman correlation Test H_o_: no association; Female: ρ=0.29, p=0.189; Male: ρ= -0.22, p=0.325) (Fig. 2D).

### Effect of *Wolbachia* infection

We next tested whether parental infection by the sex-modifying bacterium *Wolbachia* can explain the biased sex ratio observed for particular mating pairs. With the help of *Wolbachia*-specific loci, *wsp* and *ftsZ*, we evaluated the infection status of the parents from all mating pairs, and identified only 08 cases as positive for the infection – 04 each in random and inbreeding categories (Fig. S3; Supplemental data). Notably, only female parents were found infected in all the 08 cases (Fig. 3A). The infections were also limited to batch2 samples only (Fig. S3). By comparing the offspring sex ratio between the infected and uninfected female parents from the batch2, we observed that infected female parents have significantly higher proportion of female offspring, ca. 2.06 times, compared to the uninfected female parents (Wilcox test: w=24, p=0.015) (Fig. 3A). However, for 03 out of 08 infected parents, the offspring female proportions were close to 0.5 or less (Fig. 2B), suggesting that the effect of *Wolbachia*, if any, is relatively weak. Notably, the pupal mortality in *Wolbachia* infected lines were limited to maximum ∼50% of pre-pupae individuals while it extends up to ∼62% in uninfected lines (Fig. 3B).

**Figure 3.**
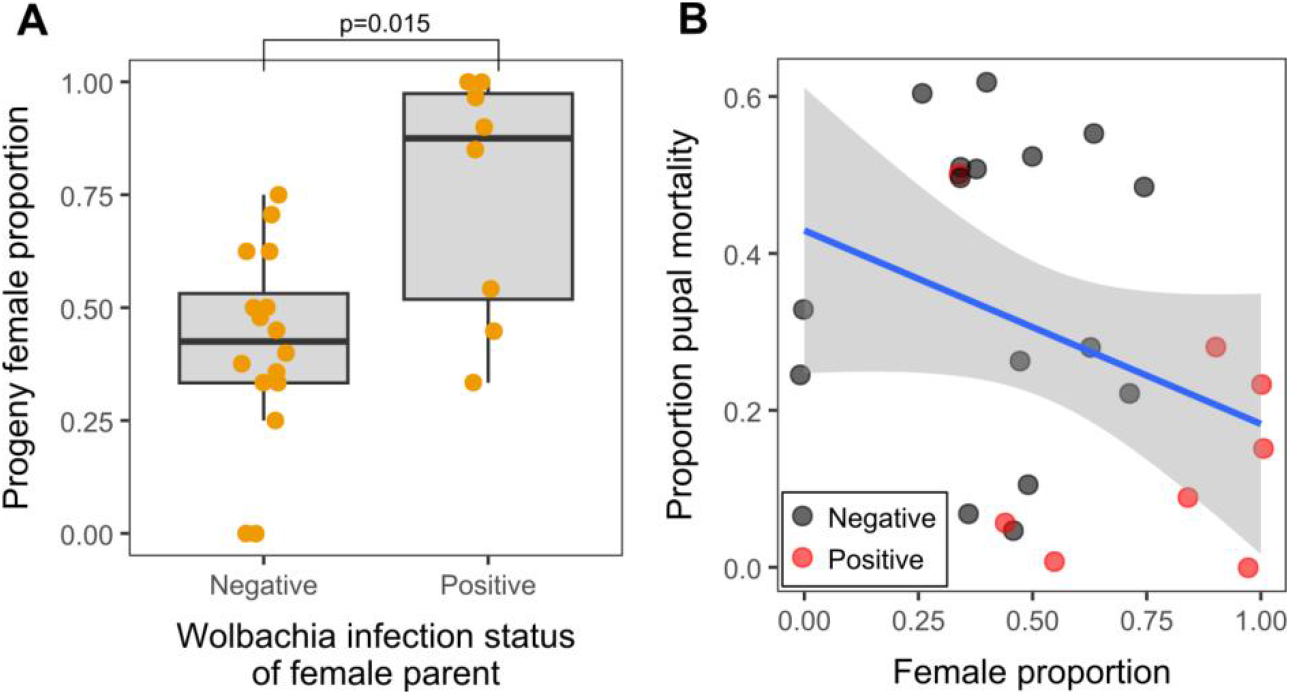
Effects of maternal infection status on sex ratio and its association with pupal mortality. (A) Box plots showing the proportion of female offspring for mating pairs differing in the infection status of the female parent (negative or positive). Male parents were uninfected in all crosses. Points represent individual mating pairs and include Batch 2 samples only (see Fig. S3 for analysis of the full dataset). Statistical comparison between groups is indicated with the corresponding p-value. (B) Scatter plot illustrating the relationship between female proportion and pupal mortality, shown separately for infected and uninfected mating lines in Batch 2. Points represent individual mating pairs. Solid lines represent linear model fits (R function ‘lm’), with shaded areas indicating 95% confidence intervals.

### *Wolbachia* strain characterization

Multi-locus sequence typing (MLST) confirmed that the *Wolbachia* from all five beetle individuals are of the same strain and identified this strain as a novel type in the PubMLST database (last checked on 27 April 2026). In the database, the closest match of this strain is ST-480, with the difference at *ftsZ* locus only: while *ftsZ*-230 allele is part of the ST-480, our strain is characterized by *ftsZ*-75 (Table S2). Nevertheless, we noticed no record of *Wolbachia* isolate for ST-480 profile in the PubMLST database. WSP-typing assigned WSP-586 to this new strain, but with an unmatched novel HVR4 region. These allelic composition of marker loci matches exactly with the previous report of *Wolbachia* in this beetle species, where the strain is labeled as *w*Bic (Sahoo et al., 2026), confirming the presence of the same *Wolbachia* strain across beetle populations. In congruence to the previous finding (Sahoo et al., 2026), the maximum-likelihood phylogenetic analyses of concatenated MLST loci placed this *Wolbachia* strain within Supergroup-A with a strong nodal support from the bootstrap analysis (Fig. S4).

## Discussion

Inbreeding is an unintended yet frequently applied component of both classical and augmentative biological control practices (Mackauer, 1976). During classical biocontrol programs, particularly in the introduction phase, a limited number of imported individuals are often inbred under constrained conditions for a prolonged period to generate sufficient population sizes for host-specificity testing and subsequent field release (Franks et al., 2011). Similarly, in augmentative practices, rapid population expansion through mass rearing can impose varying degrees of inbreeding depending on the breeding design (Nunny, 2003). Such inbreeding is likely to have detrimental consequences, including reduced genetic diversity, demographic instability, and fitness-related declines in life-history traits (Paspati et al., 2019; Ross et al., 2019; Reed et al., 2007). Despite these potential impacts, their characterization in biocontrol agents remains limited. Here, by experimentally imposing a high degree of inbreeding over a short temporal scale, we provide insights into its effects on key life-history traits in the biocontrol beetle *Z. bicolorata*.

Cross-generational full-sibling inbreeding did not produce a consistent or statistically supported shift in offspring sex ratio across the three generations examined. Median sex ratios in inbred lineages remained close to the expected 1: 1; however, these patterns warrant further evaluation under longer-term inbreeding regimes. Notably, our findings contrast with previous reports of female-biased demographic structures in this beetle species (Sushilkumar et al., 2022; Hasan & Ansari 2016; Siddhapara et al., 2012; Jayanth & Bali 1993). While multiple factors may underlie this discrepancy, differences in the level of sampling likely contribute substantially. Whereas earlier studies inferred sex ratios at the population level, our results are based on observations at brood/individual level and with constrained reproduction regime (e.g., forced monogamy, limited time windows for mating and egg-laying). Consequently, our estimates are less influenced by the processes operating over extended temporal scales or across heterogeneous population structures. For example, progeny produced over a female’s entire lifespan may integrate temporal variation in sex allocation and survival, in contrast to early egg batches, thereby yielding a dynamic cumulative sex ratio (Mappes et al., 1996; Ross et al., 2011, 2012). In addition, population-level sex ratios may be shaped by sex-specific survival (e.g., elevated male mortality due to reduced competitive ability; Teder and Kaasik, 2023), mating asymmetries arising from unequal reproductive contributions among genotypes (e.g., Wyer et al., 2024; Omkar and Afaq, 2013 (due to body size difference); Pal et al., 2024 (due to male competitiveness)) , and post-mating sex allocation strategies (Katlev et al., 2021; Firman et al., 2017). In natural settings, these effects may be further compounded by ecological and demographic factors such as sex-biased foraging behavior (Chmielewski et al., 2023), dispersal capacity (Martínez-Pérez et al., 2022), and differences in lifespan (Sielezniew et al., 2020) , all of which can collectively skew the observed sex ratio.

Cross-generational inbreeding, however, had measurable effects on pupal mortality, particularly in the later inbreeding generations. This pattern is consistent with a delayed manifestation of inbreeding depression, whereby the cumulative expression of deleterious recessive alleles reduces viability only after successive rounds of full-sibling mating (Woldemelak, 2024). Given that these observations are based on a single experimental batch, confirmation under replicated and longer-term inbreeding regimes will be necessary to establish the robustness of this trend. In contrast, adult body weight distributions remained largely unaffected by inbreeding, suggesting that growth-related traits are comparatively less sensitive to increased homozygosity in this system. Nonetheless, the pronounced sexual dimorphism observed under random mating – where females were substantially heavier than males – was attenuated under inbreeding. This reduction in dimorphism indicates a degree of sex-specific sensitivity (Vega-Trejo et al., 2022; Teder and Tammaru, 2005), wherein phenotypic differences between sexes are diminished without substantial shifts in overall trait means. Taken together, these results indicate that inbreeding does not directly alter sex allocation but can impose fitness costs through reduced survival in later generations, with effects that are trait-specific. If such fitness costs are sex-biased – for example, if one sex experiences disproportionately higher mortality – they may indirectly contribute to shifts in population-level sex ratios.

Infection by *Wolbachia* – closely matching ST-480 and likely corresponding to the strain *w*Bic (Sahoo et al., 2026) – provides a partial explanation for the observed variation in offspring sex ratio. Infected females produced significantly higher proportions of daughters compared to uninfected counterparts, consistent with the well-established capacity of *Wolbachia* to bias host reproduction (Werren et al., 2008). However, this effect was not uniform: several infected lineages exhibited near-equal or even male-biased sex ratios, indicating incomplete penetrance or context-dependent expression of the phenotype. Notably, pupal mortality in infected lineages remained comparatively lower than in some uninfected lines, suggesting that the symbiont does not impose an additional viability cost at this developmental stage and may, under certain conditions, be associated with enhanced survival. This decoupling of sex-ratio distortion and viability effects further supports the idea that *Wolbachia*-mediated phenotypes are modulated by host genetic background or environmental context (Strunov et al., 2022; Kyritsis et al., 2019; Mouton et al., 2007). Taken together, these results indicate that the sex ratio patterns are best explained by an interplay between symbiont effects, host genetic factors, and demographic or ecological processes.

## Author Contributions

BS: Methodology, Data collection, and Writing – review and editing.

RKS: Conceptualization, Data curation, Formal analysis, Funding acquisition, Investigation, Methodology, Project administration, Resources, Supervision, Validation, Visualization, Writing – original draft, and Writing – review and editing.

## Acknowledgements

This work was supported by the DST-INSPIRE Faculty Fellowship to RKS (DST/INSPIRE/04/2019/000478). We thank Karthikeyan Vasudevan for valuable academic support during the execution of this work.

## Conflicts of Interest

The authors declare no conflicts of interest.

## Data availability

Data is provided in the supplemental file. Sequences generated during the study are submitted in GenBank. Accession numbers would be available during review, if required, or upon approval for publication.

## Notes

### Competing Interest Statement

The authors have declared no competing interest.

