## Supplementary material for "Effects of cross-generational inbreeding and *Wolbachia* infection on sex ratio and life-history traits in parthenium beetle": Tables S1-S2 and Figures S1-S4

**Supplemental file** for the title, “Effects of cross-generational inbreeding and *Wolbachia* infection on sex ratio and life-history traits in parthenium beetle”.

**Table S1.** Summary statistics of sex ratio across mating categories. Median sex ratio values are presented for the random mating category and three successive generations of inbreeding (G1–G3). Median absolute deviation (MAD) and robust coefficient of variation (RCV) are provided in parentheses. Results for Batch 1 and Batch 2 are shown separately. RCV values are highlighted in bold.

| | Random | G1 | G2 | G3 |
|---|---|---|---|---|
| Batch 1 | 0.542<br>(0.055, **0.149**) | 0.612<br>(0.058, **0.141**) | 0.5<br>(0.318, **0.942**) | NAN |
| Batch 2 | 0.778<br>(0.304, **0.579**) | 0.412<br>(0.347, **1.246**) | 0.333<br>(0.17, **0.758**) | 0.5<br>(0.185, **0.549**) |

**Table S2.** MLST and WSP-based sequence typing of the strain using the PubMLST database for *Wolbachia* spp. The closest MLST allelic matches (*) differ by 1–4 nucleotides per allele. The database was last accessed on April 27, 2026.

| Sample code | **MLST** | | | | | **WSP** | | | |
|---|---|---|---|---|---|---|---|---|---|
| | coxA | fbpA | ftsZ | gatB | hcpA | HVR1 | HVR2 | HVR3 | HVR4 |
| HD578 | 33* | 36* | 75* | 32* | 42* | 52 | 57 | 39 | 255* |
| HD580 | 33* | 36* | 75* | 32* | 42* | 52 | 57 | 39 | 255* |
| HD592 | 33* | 36* | 75* | 32* | 42* | 52 | 57* | 39 | 255* |
| HDX48 | 33* | 36* | 75* | 32* | 42* | 52 | 57 | 39 | 255* |
| HDX57 | 33* | 36* | 75* | 32* | 42* | 52 | 57 | 39 | 255* |

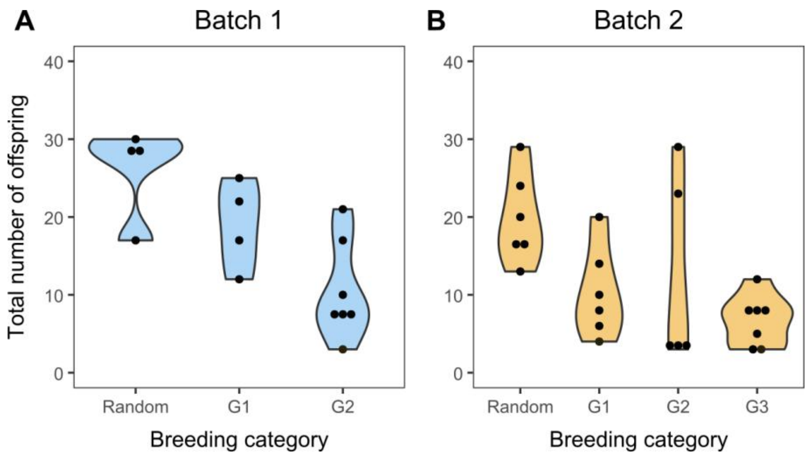


**Figure S1.** Distribution of mating lines and offspring counts across breeding categories. Violin plots show the number of mating lines and the total number of offspring per line across breeding categories. Results are presented separately for Batch 1 (A) and Batch 2 (B).

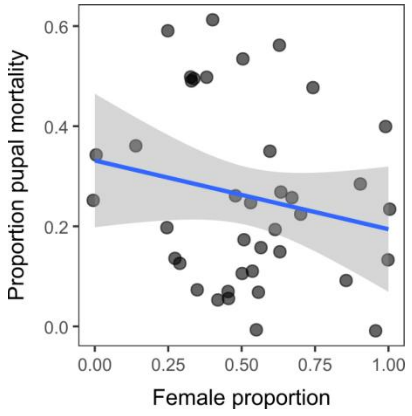


**Figure S2.** Relationship between female proportion and pupal mortality. Scatter plot illustrating the relationship between female proportion and pupal mortality using combined data from Batch 1 and Batch 2. Points represent individual mating pairs. Solid lines represent linear model fits (R function lm), with shaded areas indicating 95% confidence intervals.

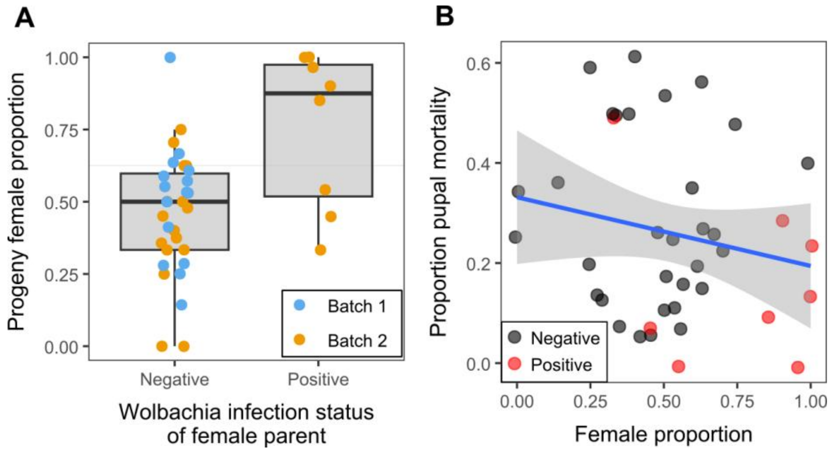


**Figure S3.** Effects of maternal infection status on sex ratio and its relationship with pupal mortality. (A) Box plots showing the proportion of female offspring for mating pairs differing in the infection status of the female parent (negative or positive). Male parents were uninfected in all crosses. Points represent individual mating pairs and include data from both Batch 1 and Batch 2. (B) Scatter plot illustrating the relationship between female proportion and pupal mortality, shown separately for infected and uninfected mating lines using combined data from Batch 1 and Batch 2. Points represent individual mating pairs. Solid lines represent linear model fits (R function lm), with shaded areas indicating 95% confidence intervals.

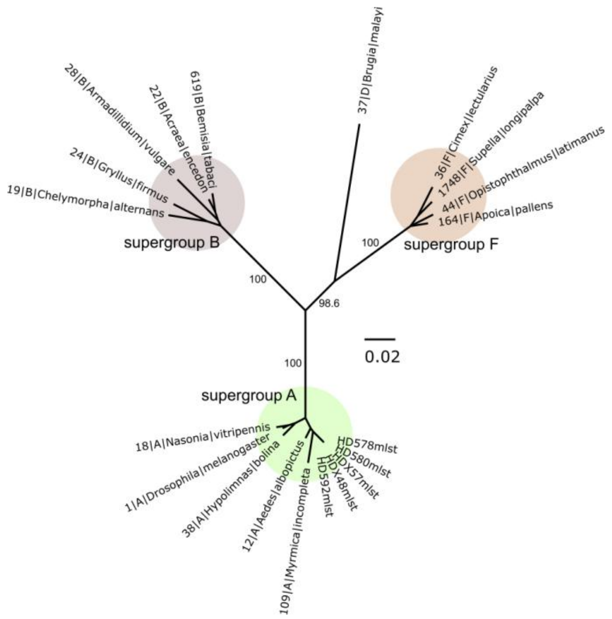


**Figure S4.** Maximum likelihood (ML) unrooted phylogenetic tree inferred from concatenated MLST sequence alignments using the Tamura-Nei (G+I) model in MEGA. Node support values were estimated from 1000 bootstrap replicates and are shown for major nodes. Samples from the present study are labeled 'mlst'.
